# Highly pathogenic avian influenza A virus (HPAIV) H5N1 infection in two European grey seals (*Halichoerus grypus*) with encephalitis

**DOI:** 10.1101/2023.05.30.542941

**Authors:** Monica Mirolo, Anne Pohlmann, Ann Kathrin Ahrens, Bianca Kühl, Ana Rubio-Garcìa, Katharina Kramer, Ulrike Meinfelder, Tanja Rosenberger, Hannah Leah Morito, Martin Beer, Martin Ludlow, Peter Wohlsein, Wolfgang Baumgärtner, Timm Harder, Albert Osterhaus

## Abstract

Recent reports documenting sporadic infections in carnivorous mammals worldwide with highly pathogenic avian influenza virus (HPAIV) H5N1 clade 2.3.4.4b have raised concerns about the potential risk of adaptation to sustained transmission in mammals, including humans. We report H5N1 clade 2.3.4.4b infection of two grey seals (*Halichoerus grypus*) from coastal waters of The Netherlands and Germany in December 2022 and February 2023, respectively. Histological and immunohistochemical investigations showed in both animals a non-suppurative and necrotizing encephalitis with viral antigen restricted to the neuroparenchyma. Whole genome sequencing showed the presence of HPAIV H5N1 clade 2.3.4.4b strains in brain tissue, which were closely related to sympatric avian influenza viruses. Viral RNA was also detected in the lung of the seal from Germany by real-time quantitative PCR. No other organs tested positive. The mammalian adaptation PB2-E627K mutation was identified in approximately 40% of the virus population present in the brain tissue of the German seal. Retrospective screening for nucleoprotein specific antibodies, of sera collected from 251 seals sampled in this region from 2020 to 2023, did not show evidence of influenza A virus specific antibodies. Similarly, screening by reverse transcription PCR of lung and brain tissue of 101 seals that had died along the Dutch coast in the period 2020-2021, did not show evidence of influenza virus infection. Collectively, these results indicate that individual seals are sporadically infected with HPAIV-H5N1 clade 2.3.4.4b, resulting in an encephalitis in the absence of a systemic infection, and with no evidence thus far of onward spread between seals.

## Introduction

Avian influenza viruses (AIVs) display a high level of diversity with respect to surface glycoproteins hemagglutinin (HA 1-16) and neuraminidase (NA 1-9) subtypes and are endemic among many species of wild birds including *Anatidae* (i.e. ducks, geese, swans), *Scolopacidae* (shorebirds/waders), and *Laridae* (gulls, terns) [1]. The AIV/H5 and AIV/H7 subtypes have been classified as low pathogenic avian influenza virus (LPAIV) or highly pathogenic avian influenza virus (HPAIV), due to differences in the usage of host-proteases involved in HA cleavage and henceforth the severity of the resulting clinical disease. Activation of the H5 and H7 of LPAIV strains is mediated by proteases expressed in the gastrointestinal tract and respiratory tract of wild birds and domestic poultry in which a mild disease is induced, including ruffled feathers and/or a drop in egg production [2]. However, AIVs encoding H5 or H7 can evolve into the HPAIV phenotype upon infection of domestic poultry, by the acquisition of multiple basic amino acids at the HA proteolytic cleavage site. This enables cleavage of the glycoprotein by the ubiquitous host protease furin, resulting in increased viral tropism and disease severity [2-4]. Consequently, HPAIV may cause severe disease in poultry and wild birds that affects multiple organs resulting in 90%-100% mortality within a few days of infection. However, HPAIV may cause milder symptoms in some waterfowl species, thus allowing increased virus to spread over long distances [3,5]. Transmission of HPAIV between birds occurs via direct or indirect contact with virus-infected secreta or excreta or by predation.

Multiple reassortment events of the HPAIV A/Goose/Guangdong/1/96 (Gs/GD H5N1) from Guangdong (China, 1996) [4] with other AIVs have resulted in ten different phylogenetic H5 clades (clades 0–9) [6]. Clade 2 has been further divided into five subgroups (clades 2.1–2.5) [6], and additional third and fourth order clade subdivisions e.g. clades 2.3.4.4a to 2.3.4.4h. The Eurasian HPAIV H5N1 bearing HA clade 2.3.4.4b was first identified in Europe in October 2020, as a reassorted Gs/GD H5NX genotype [7,8] and was subsequently identified in wild bird populations throughout Europe, Africa, Asia, and the USA. Alongside multicontinental outbreaks of Gs/GD HPAIV H5N1 infection among domestic poultry and wild birds, sporadic infections with HPAIV H5N1 were detected in domestic and free-ranging terrestrial and aquatic carnivorous mammalian species [9]. Isolated cases of HPAIV H5 clade 2.3.4.4b infection of mammals were reported in 2021 and 2022 during the ongoing epizootic in birds in Eurasia and the Americas. Sporadic infections of scavenging and predatory terrestrial mammals included the red fox (*Vulpes vulpes*), polecat (*Mustela putorius*) and badger (*Meles meles*) in The Netherlands [10]. An outbreak of HPAIV H5 clade 2.3.4.4b in farmed American minks (*Neovison vison*) in Spain in October 2022, along with indications of virus transmission between animals, has increased concerns about possible adaptation of this virus to mammals, including humans [11,12]. More recently, infection of several other species, including skunk (*Mephitis mephitis*), raccoon (*Procyon lotor*), bobcat (*Lynx rufus*), opossum (*Didelphis virginiana*), coyote (*Canis latrans*) and fisher (*Pekania pennanti*) were reported in the USA [13]. Aquatic carnivore species have also been reported to be susceptible to HPAIV H5N1 infection, including harbor seals (*Phoca vitulina*), grey (*Halichoerus grypus*) seals, porpoises (*Phocoena phocoena*), common dolphins (*Delphinus delphis*), and otters (*Lutra lutra*) [9,14,15]. Unusual mass mortality events caused by epizootics of HPAIV H5N1 infection have been reported in harbour and grey seals in North America [16], and sea lions (*Otaria flavescens* and *Arctocephalus australis*) in South America [17]. Analysis of HPAIV H5N1 strains identified in mammals has shown the presence of mutations indicative for adaptation to mammals [9]. This includes E627K in PB2, which enhances virus replication in mammalian cells, while reducing its capacity to replicate in avian cells [18]. Such mammalian adaptive mutations can serve as molecular markers to facilitate assessment of the pandemic risk posed by circulating AIVs.

In this study, we combine data about the identification and characterization of HPAIV H5N1 2.3.4.4b in two grey seals from the Dutch and German North Sea seal population, referred to as case 1 and 2, that have initially been individually studied in two German research institutes. Both seals died with an encephalitis, and the virus was recovered from the brain of both seals and in the lung of case 2. Retrospective serological and molecular screening of hundreds of grey and harbour seal samples did not provide further evidence of AIV infections among seals in this region in the past three years.

## Materials and Methods

### Clinical samples

Case 1. The Sealcentre Pieterburen, Pieterburen, The Netherlands, provided clinical samples (lung, brain, spleen, kidney, and liver tissue and oropharyngeal swabs) from an adult male grey seal (approximately eight years-old), which had died on the beach of the Dutch island of Terschelling (NL) in December 2022, with respiratory and neurological clinical signs. These included laboured breathing, ataxia and body tremors. Following natural death, the seal was transported to the Sealcentre Pieterburen located in Pieterburen, The Netherlands, for gross pathology analysis. Collection of dead seals for diagnostic purposes by the Sealcentre Pieterburen is permitted by the government of The Netherlands (application number FF/ 75/2012/015). Necropsy was performed at the Sealcentre Pieterburen in Pieterburen (NL) under stringent protective measures, and clinical samples were stored at −80°C for virological and molecular analyses and additional tissues were fixed in 10% buffered formalin for patho-histological analyses. Samples were transported to the Research Center for Emerging Infections and Zoonoses, University of Veterinary Medicine Hannover, Germany for further analysis under permit number DE 03 201 0043 21 obtained from the Fachbereich Öffentliche Ordnung, Gewerbe-und Veterinärangelegenheiten, Hannover, Germany. Serum samples were collected from seals at the Sealcentre Pieterburen for routine diagnostic procedures during rehabilitation of the animals.

Case 2. A grey seal pup (approximately 2 weeks-old) that was admitted to the Seal Rehabilitation Centre at Friedrichskoog, on the Wadden Sea coast of Germany on January 1st, 2023, developed well until February 18th, when it became anorectic and showed reluctance to dive. It died February 26^th^ without further conspicuous clinical signs. Necropsy was performed one day after death at the Landeslabor Schleswig-Holstein (LSH), Neumünster, Germany following the LSH standard procedure for seals. Gross pathology analysis and sampling of the animal was conducted under ethical permit of the Ministry of Energy, Agriculture, Environment and Rural Areas of Schleswig-Holstein, Germany (Permit number V 312-72241.121-19 (70-6/07)). No further seals had been admitted since the February 4th. The other seals present in the centre stayed healthy until being released in April. Swab samples were obtained twice from 19 grey seals and 1 harbour seal at the centre and additional blood samples were taken. Tissue samples for PCR analysis were taken and directly analysed. Furthermore, tissues for patho-histological analyses were fixed in 10% buffered formalin and parasitological investigation on intestinal content were performed.

### Histology and immunohistochemistry

The fixed tissue samples from case 1 and 2 were routinely embedded in paraffin wax, sectioned at 3 µm and stained with haematoxylin and eosin. Immunohistochemistry (IHC) was performed on paraffin-embedded tissue sections using a murine monoclonal antibody specific for the nucleoprotein (CLONEGENE LLC, HB65) [19] according to a previously described procedure [20]. Immunohistochemistry for morbillivirus antigen was performed as described previously [21].

### Virus detection and subtyping

Frozen tissues (approximately 60 mg) obtained at the necropsy were homogenized in 500 µl in phosphate-buffered saline (PBS) solution using ceramic beads in a FastPrep-24 5G homogenizer (MP Biomedical), and then centrifuged at 12,000 RCF for 5 minutes. RNA isolation from 140 µl of this clarified supernatant from tissues was performed using a QIAamp Viral RNA Mini Kit protocol (Qiagen). Subsequently, RNA extracted from multiple organs was tested by a reverse transcription PCR (RT-PCR) or by real time quantitative PCR (RT-qPCR) using primers already available in each institute for detection of phocid alphaherpes virus 1 (PhHV-1), phocine or canine distemper virus (PDV, CDV, *Canine, Phocine morbillivirus*). The primer sets that were used have been previously designed to detect the presence of herpesviruses and paramyxoviruses, and CDV [22-24]. The amplified RT-PCR amplicons were analysed by gel electrophoresis, purified using the Monarch^®^ DNA Gel Extraction Kit (New England BioLabs), Sanger sequenced, and analysed using BLAST (Basic Local Alignment Search Tool) with the GenBank NCBI nucleotide database. A RT-qPCR targeting a conserved region of the AIV matrix gene (MP RT-qPCR) was used to test all clinical samples of case 1 [25]. RNA from multiple organs of seal from case 2 was also assayed by a specific RT-qPCR to detect and subtype the virus from case 2 [26]. AIV subtyping of case 1 was instead performed using specific HA and NA primers (Table S1).

### Virus isolation

To prove the viral etiology of the encephalitis and to investigate the acquisition of genetic signatures of mammalian adaptation in the avian H5N1 virus upon infection of the new host [18] the H5N1 virus from case 1 was cultured on mammalian cells. For this purpose, Madin-Darby Canine Kidney (MDCK) cells were cultivated in Dulbecco’s Modified Eagle Medium (DMEM) supplemented with 10% fetal bovine serum and penicillin (100 IU/mL)/streptomycin (100 µg/mL) (1% Pen/strep). Then, 100 µl of clarified supernatant from homogenized brain tissue of the seal from case 1 was inoculated onto an 80% confluent MDCK monolayer cultivated in DMEM supplemented with 0.1% BSA, 1% Pen/strep, 1% Glutamax, 1 × MEM non-essential amino acids solution, 1M HEPES and 1 µg/mL of TPCK in T25 tissue culture flasks.

RNA was extracted from supernatant of inoculated cultures at 0-and 3-days post inoculation (dpi) and virus replication was assessed using an MP RT-qPCR. Full-length H5N1 genomes from cell supernatant collected at 2 dpi and 3 dpi were assayed by deep sequencing together with the original clinical samples.

### Genome sequencing

Full-genome sequencing of original case 1 and case 2 clinical samples and MDCK isolates (2 and 3 dpi) was performed using a previously described nanopore-based real-time sequencing method with prior full genome amplification [27]. In brief, RNA was extracted using TRIZOL and RNeasy Mini Kit (Qiagen, Germany) and genome amplification was performed via a universal AIV-End-RT-PCR using Superscript III One-Step and Platinum Taq (Thermo Fisher Scientific, USA). This used one primerpair (Pan-IVA-1F: TCCCAGTCACGACGTCGTAGCGAAAGCAGG; Pan-IVA-1R: GGAAACA GCTATGACCATGAGTAGAAACAAGG), which binds to the conserved ends of the AIV genome segments. PCR products were purified with AMPure XP Magnetic Beads (Beckman-Coulter, USA), prior to full-genome sequencing on a MinION platform (Oxford Nanopore Technologies, ONT, UK) using Rapid Barcoding Kit (SQK-RBK114-24 or SQK-RBK004, ONT) for transposon-based library preparation and multiplexing. Sequencing was performed according to the manufacturer’s instructions with R10.4.1 or R9.4.1 flow cells on an Mk1b device with MinKNOW Software Core (v5.4.3). Live high accuracy base calling of the raw data with Guppy (v6.4.4, ONT) was followed by demultiplexing, a quality check and a trimming step to remove low quality, primer and short (<20 bp) sequences. Full-length genome consensus sequences were generated in a map-to-reference approach utilizing MiniMap2 [28]. A curated collection of all HA and NA subtypes alongside an assortment of internal gene sequences were chosen as reference genomes to cover all potentially circulating viral strains. Polishing of the final genome sequences was performed manually after consensus production according to the highest quality (60%) in Geneious Prime (v2021.0.1, Biomatters, New Zealand).

### Phylogenetic analyses

Segment specific and concatenated whole-genome multiple alignments were generated for both genomes with H5N1 genomes collected worldwide since October 2020 using MAFFT (v7.450) [29] and subsequent maximum likelihood (ML) trees were calculated with RAxML (v8.2.11) [30] utilizing model GTR GAMMA with rapid bootstrapping and search for the best scoring ML tree supported with 1000 bootstrap replicates or alternatively with FastTree (v2.1.11) [31]. Subsets of close relative genomes were extracted and used for further ML phylogenetic analyses. Each segment was aligned with homologous segments from closely related avian strains using Molecular Evolutionary Genetics Analysis (MEGA) 11 [32].

### Serological analysis

Serum samples collected routinely post-mortem from the two diseased seals and from convalescent gr*e*y seals and harbor seals in The Netherlands (n= 325) and in Germany (n= 20) from 2020 to 2023, were tested retrospectively with a commercial nucleoprotein specific competitive ELISA (ID VET, ID Screen Influenza A Antibody Competition Multi-species). The sera were heat-inactivated at 56°C for 30 min and diluted 1:20 in PBS solution prior to testing.

## Results

### Detection and isolation of AIV from grey seals

Case 1. Macroscopic finding at necropsy included dark-red coloration of left/right lung. No ectoparasites or liver-, heart-, and lungworms were observed. A multifocal moderate acute, lympho-histiocytic and necrotizing encephalitis with mild gliosis and neuronal necroses was observed upon histological analysis (Figure 1A). A focally mild, lympho-histiocytic pneumonia was present in the lung tissue. Intralesional nucleoprotein antigen of AIV was detected only in the brain of this animal (Figure 1B). Given the observation of neurological clinical signs, initial RT-PCR testing of RNA extracted from brain tissue was performed for neurotropic viruses, which have been previously documented in seal populations in the North Sea. No amplicons were detected for PhHV-1 and CDV/PDV in the brain (data not shown). Morbillivirus antigen was not detected by IHC of brain sections from this animal. Testing of RNA extracted from lung, brain, liver, spleen, and kidney tissue and throat/nose swabs for AIV using a MP RT-qPCR showed a positive signal in brain tissue (Ct, 16.60).

**Figure 1.**
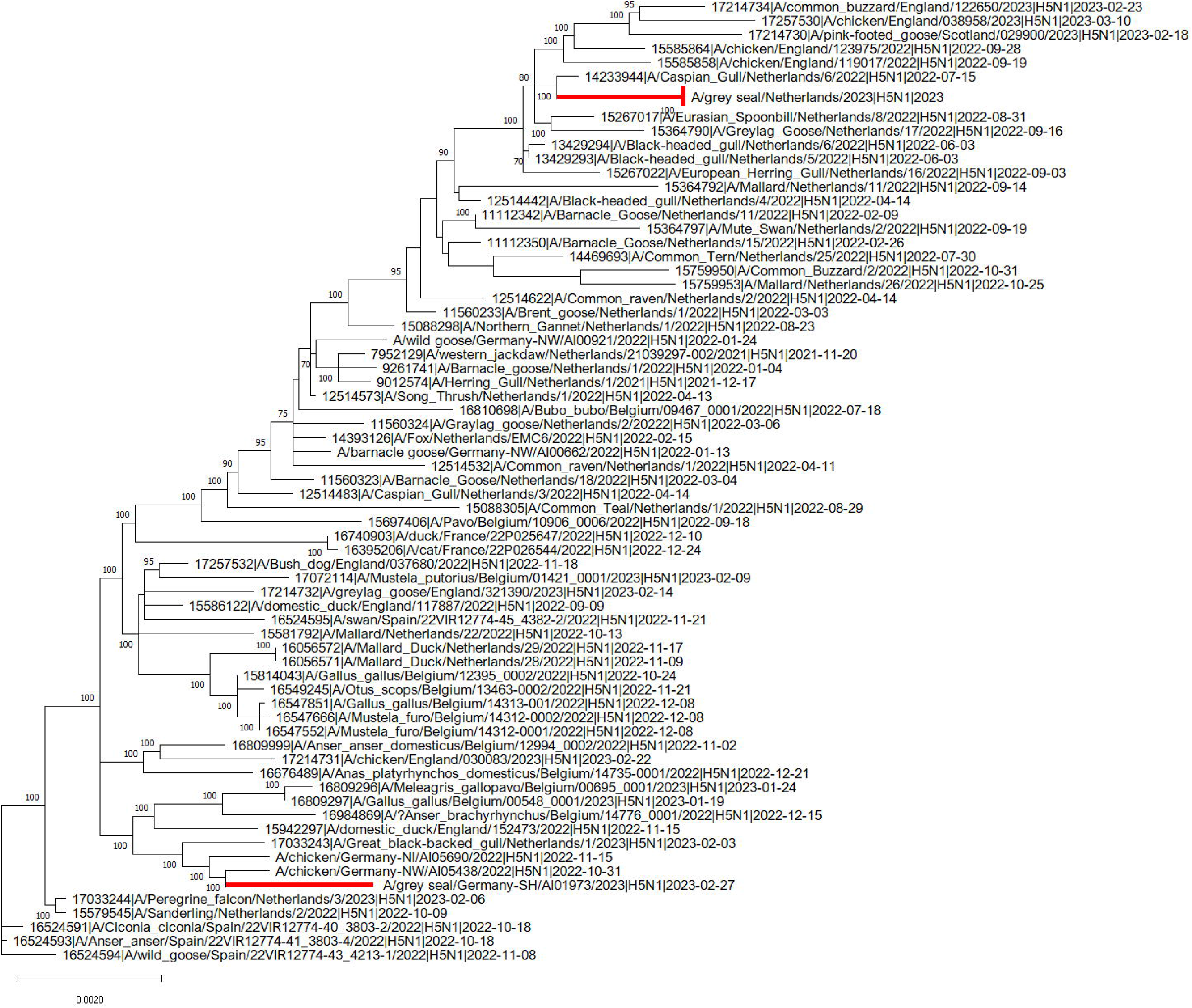
Histopathology and immunohistochemical findings in the brain tissue of the seals. The cerebellum of the H5N1 infected seal from The Netherlands (case 1) showed the following changes: A. An acute non-suppurative and necrotizing encephalitis with mild perivascular lympho-histiocytic infiltration (circle), diffuse gliosis of the adjacent neuroparenchyma (molecular layer), necrosis of Purkinje cells (arrow) and necrosis of inner granular cells (arrowheads); B. Abundant intralesional AIV nucleoprotein in cells of the inner granular layer (arrowheads) and within the adjacent neuropil. The brain of the H5N1 infected seal from Germany (case 2) showed: C. Moderate lympho-histiocytic leptomeningitis. D. Moderate non-suppurative and necrotizing encephalitis with perivascular lympho-histiocytic cuffing and vasculitis (arrows), gliosis and necrosis (arrowheads) in the adjacent parenchyma. E. Abundant AIV nucleoprotein in neuronal nuclei, perikarya and processes (arrows) and in various glial cells (arrowheads) of the cerebrum.

Case 2. Macroscopic observation at necropsy showed that the seal pup was in good body condition. The lung showed failure to collapse, diffuse firmness and patchy dark red parenchyma with hyperplastic lymph nodes. In addition, moderate thymic atrophy, fibrinous perisplenitis and perihepatitis and catarrhal enteritis was obvious. The brain was macroscopically without pathological findings. No ecto- or endoparasites were found macroscopically. In addition, endoparasites were not detected, upon microscopic investigation of the intestinal content. The dead seal displayed a multifocal to coalescing moderate to severe lympho-histiocytic meningoencephalitis lymphohistiocytic to necrotizing vasculitis and and widespread single cell necroses in the neuroparenchyma (Figure 1 C, D). In the lung, a multifocal mild to moderate non-suppurative interstitial pneumonia with lymphohistiocytic vasculitis was found.

Depletion of lymphocytic organs and vasculitis in several organs were found additionally. Quantitative PCRs (qPCR) specific for CDV (in lunge tissue) were negative. However, qPCR for AIV was positive in lung and brain tissue. Lung tissue was used for further specification as AIV H5. IHC revealed viral antigen only in the brain located in the nuclei, perikarya and processes of neurons as well as in nuclei and cytoplasms of glial cells (Figure 1E).

### Molecular and phylogenetic analysis

Following molecular confirmation of AIV infection in case 1 and case 2, the specific subtype was identified using H5 and N1 specific RT-PCRs, given the recent detection of H5N1 AIV in carnivore species in The Netherlands [10]. Sanger sequencing of positive amplicons showed the presence of the PLRE-KRRKR multiple basic cleavage site in the HA, which classified these strains within the HPAIV H5 pathotype. The complete consensus genome sequence of both strains was obtained by next generation sequencing to enable assessment of the evolutionary relationship between the strains and identify possible mutations indicative of mammalian adaptation. Analysis of a phylogenetic tree based on concatenated whole genome sequences showed that the HPAIV H5N1 strains from case 1 and case 2 are closely related to sympatric wild and domestic bird H5N1 isolates of H5 from clade 2.3.4.4b, but cluster into geographically distinct clades (Figure 2). Both genomes were H5N1 reassortants (Ger-10-21-N1.5/Rus-09-21-N1; A/duck/Saratov/29-02/2021-like) dominating the H5N1 cases in Europe 2023 and circulating in Germany and The Netherlands since November 2021 [33].

**Figure 2:**
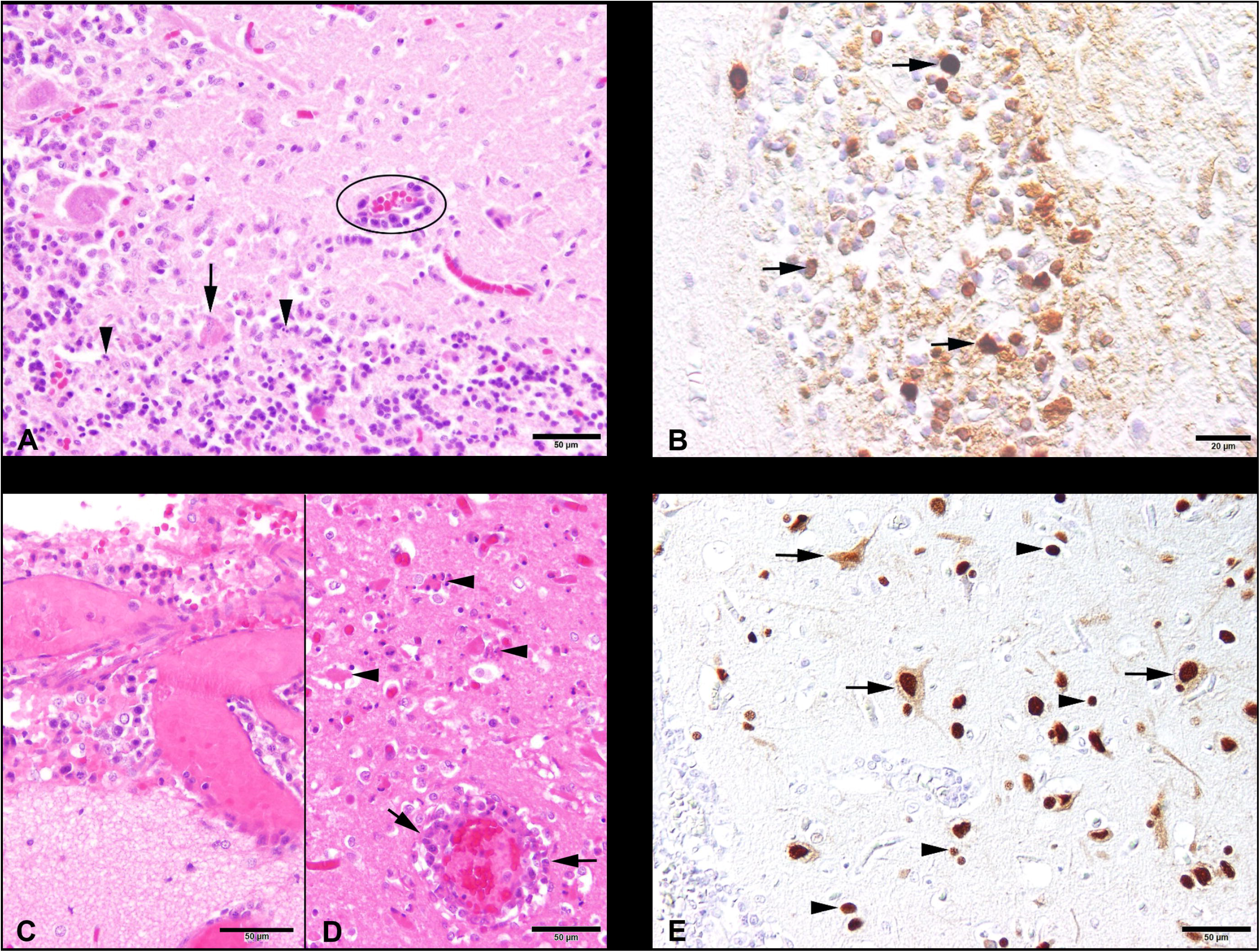
Phylogenetic relationships of concatenated consensus whole-HPAIV H5N1 sequences from The Netherlands and Germany in 2022-2023. The complete genome sequence of H5N1 strains derived from infected seals in Germany and The Netherlands were compared to sympatric avian strains belonging to the clade 2.3.4.4b of H5. Sequences were obtained from the EpiFluTM GISAID database. Isolate identifier contains the following information: Accession number, virus designation, subtype and collection date. Maximum likelihood (ML) trees were calculated with RAxML (v8.2.11) utilizing model GTR GAMMA with rapid bootstrapping. Scale bar indicates nucleotide substitutions per site. Bootstrap values below 70 are not displayed.

Neither of the seal HPAIV strains possessed amino acid substitutions in the HA protein such as Q226L and G228S, which are known to enhance binding of the H5 viruses to human-like receptor α-2,6 [34]. However, approximately 40% of the virus population present in the brain of case 2 from Germany displayed the well-characterized E627K mutation in PB2, whereas 100% of the virus population in tissue of case 1 from The Netherlands displayed the avian 627E variant in PB2.

### Virus isolation

AIV was isolated in MDCK cells from supernatant of homogenised and centrifuged brain tissue supernatant in the Hannover BSL3 setting. Extensive cytopathogenic effects including cell rounding, ballooning, and vacuolization were visible at 3 dpi. Assessment of virus replication by MP RT-qPCR showed a reduction in Ct values from 33 to 11 between day 0 and day 3 post-inoculation. No genetic mutations have been detected in the virus at 2 and 3 dpi compared to the original virus from the brain.

### Retrospective screening of seals for AIV infections (2020-2023)

Retrospective screening of sera collected from grey (n= 59) and harbour seals (n = 266) sampled in the same geographic area from 2020 to 2023 did not show AIV specific antibodies in a competitive multi-species ELISA. Similarly, screening by RT-PCR of lungs and brains of 101 grey seals (n= 12) and harbour seals (n= 89) that had died in the same area between years 2020-2021, did not provide evidence of AIV infection. A total of 20 serum samples taken from 19 further grey seals pups and one harbour seals 14 days after the death of the grey seal pup in the Friedrichskoog centre tested negative for influenza A virus antibodies indicating that no further spill-over infections had occurred at the centre.

## Discussion

Since 2021, the HPAIV H5N1 epizootic has been associated with multiple AIV outbreaks in Europe and beyond, causing mass mortalities amongst wild birds, and domestic poultry, the latter of which has necessitated targeted culling of infected flocks and associated movement restrictions. In this study, we describe HPAIV H5N1 clade 2.3.4.4b infection of two grey seals originating from Dutch and German coastal areas, respectively. Case 1 have been infected in the wild, whereas case 2 became infected several weeks after its admission to the German rehabilitation centre, most likely through indirect contacts with gulls that frequented the centre during feeding times. The clinical manifestations of case 1 were indicative of central nervous system (CNS) disease. Alterations in the lungs of both animals were observed by gross pathological examination interstitial pneumonia was confirmed by histological examination in case 2. Extensive histological lesions indicative of an acute encephalitis, with detection of AIV antigen within these lesions by IHC and positive MP-RT-qPCR of the brains of both cases identified the viral cause of death of both animals. Other known viral causes of encephalitis in seals such as herpesvirus and morbillivirus infections [35-38] could not be demonstrated by molecular analyses of brain tissue from case 1 and case 2. Finally, the isolation of HPAIV H5N1 from the brain of case 1, confirmed that this virus was the cause of the clinical disease and the observed histological lesions.

It remains unclear how HPAIV H5N1 reached the brain of seals in the absence of detectable viremia or systemic virus replication. It has been suggested that H5N1 may spread to the CNS via infection of olfactory nerves in the nasal cavity with subsequent axonal transport leading to infection of the olfactory bulb, or alternatively through the myenteric plexus from the intestine [39,40]. Similarly, individual cases of encephalitis caused by HPAIV H5N8 infection have been reported from the same geographical region [41]. No indications were found for horizontal spread of HPAIV H5N1 detected in case1 and case 2. This may be related to sudden death due to viral encephalitis. However, analysis of previous outbreaks of AIV H10N7 in seals in the North Sea has shown that horizontal transmission of AIV among seals is possible [42,43]. Similarly, probable horizontal transmission of HPAIV H5N1 has been reported following infection of farmed minks in Spain, in which high numbers of animals were co-housed [11]. Although transmission of HPAIV H5N1 among seals has not yet been confirmed, seal-to-seal transmission could not be excluded among four seals during the recent outbreak in seals on the New England coast, due to the sequence similarity of the virus strains [16]. Alternatively, these animals may have been infected by the same source, most likely dead or moribund infected bird or bird excreta, given that seals share the coastal habitat with multiple bird species [17,41]. Combined retrospective serological screening of seal sera and molecular analysis of lung and brain tissues of dead animals as well as a previous serological study (40), showed no evidence of further AIV infections along the North Sea coast of The Netherlands and Germany between 2020 and 2022.

The HPAIV H5N1 infected of case 2 displayed a minority virus population (approximately 40%) in the brain with the PB2-E627K mutation. This probably arose upon host switching from bird to mammal [18]. Other mutations such as Q226L and G228S in HA which induce a shift from avian receptor specificity to the α-2,6 human-like receptors, were not observed [44]. Thus far H5N1 clade 2.3.4.4b strains have caused sporadic infections in humans worldwide, including individual cases in China, UK, Vietnam, USA, and two cases in Spain, all of them showing symptoms of a mild febrile illness [45]. Since other AIVs, including HPAIV H5N1 of other clades have caused fatal infections in humans, extensive monitoring of AIVs in birds and mammals is of the utmost importance. Furthermore, human contacts with HPAIV H5N1 infected animals and their excreta should be avoided to mitigate the risk of zoonotic infections.

## Conclusions

We have shown that probably upon direct or indirect contact with infected birds, HPAIV H5N1 clade 2.3.4.4b may sporadically cause fatal infections of seals, without so far evidence of seal-to-seal transmission. The widespread epizootic of HPAIV H5N1 clade 2.3.4.4b among wild birds in Eurasia and the Americas apparently increases the risk of infection of wild mammalian carnivore species, and consequently enhances the chance of emerging mammalian-adaption mutations. Increased surveillance and molecular characterization of HPAIV H5N1 clade 2.3.4.4b in wild birds, domestic poultry and carnivorous aquatic and terrestrial mammals is therefore of the utmost importance to better inform ongoing risk assessment of potential for sustained transmission of these viruses in mammals and eventually in humans.

## Supporting information

Suppl. table S1_Mirolo et al.,_H5N1 infection in two grey seals from European waters

## Acknowledgements

We are grateful to the staff and volunteers of the Sealcentre Pieterburen, Pieterburen, The Netherlands and to the Seehundstation Friedrichskoog, Friedrichskoog, Germany, for providing the tissues, swabs and serum samples that were analysed in the present study. We thank Dr. Alexander Postel for providing the oligonucleotides used in this study for AIV subtyping. We also acknowledge the authors, originating and submitting laboratories of the sequences obtained from GISAID’s EpiFlu Database (http://www.gisaid.org).

## References

1. Stallknecht DE, Shane SM. Host range of avian influenza virus in free-living birds. Vet Res Commun. 1988;12(2-3):125–41.

2. Franca MS, Brown JD. Influenza pathobiology and pathogenesis in avian species. Curr Top Microbiol Immunol. 2014;385:221–42.

3. Brown JD, Stallknecht DE, Swayne DE. Experimental infection of swans and geese with highly pathogenic avian influenza virus (H5N1) of Asian lineage. Emerg Infect Dis. 2008 Jan;14(1):136–42.

4. Xu X, Subbarao Cox NJ, et al. Genetic characterization of the pathogenic influenza A/Goose/Guangdong/1/96 (H5N1) virus: similarity of its hemagglutinin gene to those of H5N1 viruses from the 1997 outbreaks in Hong Kong. Virology. 1999 Aug 15;261(1):15–9.

5. Keawcharoen J, van Riel D, van Amerongen G, et al. Wild ducks as long-distance vectors of highly pathogenic avian influenza virus (H5N1). Emerg Infect Dis. 2008 Apr;14(4):600–7.

6. Group WOFHNEW. Toward a unified nomenclature system for highly pathogenic avian influenza virus (H5N1). Emerg Infect Dis. 2008 Jul;14(7):e1.

7. Authority EFS, Prevention ECfD, Control, et al. Avian influenza overview February – May 2020. EFSA Journal. 2020;18(6):e06194.

8. European Food Safety A, European Centre for Disease P, Control, et al. Avian influenza overview June -September 2022. EFSA J. 2022 Oct;20(10):e07597.

9. European Food Safety A, European Centre for Disease P, Control, et al. Avian influenza overview December 2022 - March 2023. EFSA J. 2023 Mar;21(3):e07917.

10. Vreman S, Kik M, Germeraad E, et al. Zoonotic Mutation of Highly Pathogenic Avian Influenza H5N1 Virus Identified in the Brain of Multiple Wild Carnivore Species. Pathogens. 2023 Jan 20;12(2).

11. Aguero M, Monne I, Sanchez A, et al. Highly pathogenic avian influenza A(H5N1) virus infection in farmed minks, Spain, October 2022. Euro Surveill. 2023 Jan;28(3).

12. de Vries E, de Haan CA. Letter to the editor: Highly pathogenic influenza A(H5N1) viruses in farmed mink outbreak contain a disrupted second sialic acid binding site in neuraminidase, similar to human influenza A viruses. Euro Surveill. 2023 Feb;28(7).

13. Elsmo E, Wünschmann A, Beckmen K, et al. Pathology of natural infection with highly pathogenic avian influenza virus (H5N1) clade 2.3.4.4b in wild terrestrial mammals in the United States in 2022. bioRxiv. 2023:2023.03.10.532068.

14. Puryear W, Sawatzki K, Hill N, et al. Highly Pathogenic Avian Influenza A(H5N1) Virus Outbreak in New England Seals, United States. Emerg Infect Dis. 2023 Apr;29(4):786–791.

15. Thorsson E, Zohari S, Roos A, et al. Highly Pathogenic Avian Influenza A(H5N1) Virus in a Harbor Porpoise, Sweden. Emerg Infect Dis. 2023 Apr;29(4):852–855.

16. Puryear W, Sawatzki K, Hill N, et al. Outbreak of Highly Pathogenic Avian Influenza H5N1 in New England Seals. bioRxiv. 2022:2022.07. 29.501155.

17. Leguia M, Garcia-Glaessner A, Munoz-Saavedra B, et al. Highly pathogenic avian influenza A (H5N1) in marine mammals and seabirds in Peru. bioRxiv. 2023:2023.03. 03.531008.

18. Bordes L, Vreman S, Heutink R, et al. Highly Pathogenic Avian Influenza H5N1 Virus Infections in Wild Red Foxes (Vulpes vulpes) Show Neurotropism and Adaptive Virus Mutations. Microbiol Spectr. 2023 Feb 14;11(1):e0286722.

19. de Boer GF, Back W, Osterhaus AD. An ELISA for detection of antibodies against influenza A nucleoprotein in humans and various animal species. Arch Virol. 1990;115(1-2):47–61.

20. Kuhl B, Beyerbach M, Baumgartner W, et al. Characterization of microglia/macrophage phenotypes in the spinal cord following intervertebral disc herniation. Front Vet Sci. 2022;9:942967.

21. Jo WK, Grilo ML, Wohlsein P, et al. Dolphin Morbillivirus in a Fin Whale (Balaenoptera physalus) in Denmark, 2016. J Wildl Dis. 2017 Oct;53(4):921–924.

22. VanDevanter DR, Warrener P, Bennett L, et al. Detection and analysis of diverse herpesviral species by consensus primer PCR. J Clin Microbiol. 1996 Jul;34(7):1666–71.

23. Tong S, Chern SW, Li Y, et al. Sensitive and broadly reactive reverse transcription-PCR assays to detect novel paramyxoviruses. J Clin Microbiol. 2008 Aug;46(8):2652–8.

24. Scagliarini A, Dal Pozzo F, Gallina L, et al. TaqMan based real time PCR for the quantification of canine distemper virus. Vet Res Commun. 2007 Aug;31 Suppl 1:261–3.

25. Munster VJ, Wallensten A, Baas C, et al. Mallards and highly pathogenic avian influenza ancestral viruses, northern Europe. Emerg Infect Dis. 2005 Oct;11(10):1545–51.

26. Hassan KE, Ahrens AK, Ali A, et al. Improved Subtyping of Avian Influenza Viruses Using an RT-qPCR-Based Low Density Array: ‘Riems Influenza a Typing Array’, Version 2 (RITA-2). Viruses. 2022 Feb 17;14(2).

27. King J, Harder T, Beer M, et al. Rapid multiplex MinION nanopore sequencing workflow for Influenza A viruses. BMC Infect Dis. 2020 Sep 3;20(1):648.

28. Li H. Minimap2: pairwise alignment for nucleotide sequences. Bioinformatics. 2018 Sep 15;34(18):3094–3100.

29. Katoh K, Standley DM. MAFFT multiple sequence alignment software version 7: improvements in performance and usability. Mol Biol Evol. 2013 Apr;30(4):772–80.

30. Stamatakis A. RAxML version 8: a tool for phylogenetic analysis and post-analysis of large phylogenies. Bioinformatics. 2014 May 1;30(9):1312–3.

31. Price MN, Dehal PS, Arkin AP. FastTree: computing large minimum evolution trees with profiles instead of a distance matrix. Mol Biol Evol. 2009 Jul;26(7):1641–50.

32. Tamura K, Stecher G, Kumar S. MEGA11: Molecular Evolutionary Genetics Analysis Version 11. Mol Biol Evol. 2021 Jun 25;38(7):3022–3027.

33. Pohlmann A, King J, Fusaro A, et al. Has Epizootic Become Enzootic? Evidence for a Fundamental Change in the Infection Dynamics of Highly Pathogenic Avian Influenza in Europe, 2021. mBio. 2022 Aug 30;13(4):e0060922.

34. Gao Y, Zhang Y, Shinya K, et al. Identification of amino acids in HA and PB2 critical for the transmission of H5N1 avian influenza viruses in a mammalian host. PLoS pathogens. 2009;5(12):e1000709.

35. Baily JL, Willoughby K, Maley M, et al. Widespread neonatal infection with phocid herpesvirus 1 in free-ranging and stranded grey seals Halichoerus grypus. Dis Aquat Organ. 2019 Mar 14;133(3):181–187.

36. Gulland FM, Lowenstine LJ, Lapointe JM, et al. Herpesvirus infection in stranded Pacific harbor seals of coastal California. J Wildl Dis. 1997 Jul;33(3):450–8.

37. Jo WK, Pfankuche VM, Lehmbecker A, et al. Association of Batai Virus Infection and Encephalitis in Harbor Seals, Germany, 2016. Emerg Infect Dis. 2018 Sep;24(9):1691–1695.

38. Bodewes R, Rubio García A, Wiersma LC, et al. Novel B19-like parvovirus in the brain of a harbor seal. PLoS One. 2013;8(11):e79259.

39. Rimmelzwaan GF, van Riel D, Baars M, et al. Influenza A virus (H5N1) infection in cats causes systemic disease with potential novel routes of virus spread within and between hosts. Am J Pathol. 2006 Jan;168(1):176-83; quiz 364.

40. van Riel D, Verdijk R, Kuiken T. The olfactory nerve: a shortcut for influenza and other viral diseases into the central nervous system. J Pathol. 2015 Jan;235(2):277–87.

41. Postel A, King J, Kaiser FK, et al. Infections with highly pathogenic avian influenza A virus (HPAIV) H5N8 in harbor seals at the German North Sea coast, 2021. Emerg Microbes Infect. 2022 Dec;11(1):725–729.

42. Bodewes R, Bestebroer TM, van der Vries E, et al. Avian Influenza A(H10N7) virus-associated mass deaths among harbor seals. Emerg Infect Dis. 2015 Apr;21(4):720–2.

43. Krog JS, Hansen MS, Holm E, et al. Influenza A(H10N7) virus in dead harbor seals, Denmark. Emerg Infect Dis. 2015 Apr;21(4):684–7.

44. Sun H, Pu J, Wei Y, et al. Highly Pathogenic Avian Influenza H5N6 Viruses Exhibit Enhanced Affinity for Human Type Sialic Acid Receptor and In-Contact Transmission in Model Ferrets. J Virol. 2016 Jul 15;90(14):6235–6243.

45. Assessment RR. Assessment of risk associated with recent influenza A (H5N1) clade 2.3. 4.4 b viruses. 2022.

