## Supplementary material for "Highly pathogenic avian influenza A virus (HPAIV) H5N1 infection in two European grey seals (*Halichoerus grypus*) with encephalitis": Suppl. table S1_Mirolo et al.,_H5N1 infection in two grey seals from European waters

Table S1. AIV segment specific primers used in this study.

| **Primer** | **Sequence (5‘-3‘)** | **Target** | **Reference** |
| --- | --- | --- | --- |
| H5-start-f | GCAGGGGTTCACTCTGTCAAAATG | H5 | 1 |
| H5-mid-1035r | CTATTTCTGAGCCCAGTCGCAAGG | H5 | 1 |
| H5-mid-815f | CCAGAAWATGCATACAAAATTGTCAARAA | H5 | 2 |
| H5-uni-r | ACAAGGGTGTTTTTAACTACAATCTGAACTC | H5 | 2 |
| NA Fw (15bp) | AGCAAAAGCAGGAGT | NA | 3 |
| NA Rv (21bp) | AGTAGAAACAAGGAGTTTTTT | NA | 3 |

References:

1. Postel *et al*., 2022; doi 10.1080/22221751.2022.2043726
2. Shin *et al.*, 2019; doi 10.3201/eid2512.181472
3. Perk *et al.,* 2007; doi 10.1007/s11262-007-0120-1
